# Recent improvements to the automatic characterization and data collection algorithms on MASSIF-1

**DOI:** 10.1101/236596

**Authors:** Olof Svensson, Maciej Gilski, Didier Nurizzo, Matthew W. Bowler

## Abstract

Macromolecular crystallography (MX) is now a mature and widely used technique essential in the understanding of biology and medicine. Increases in computing power combined with robotics have enabled not only large numbers of samples to be screened and characterised but also for better decisions to be taken on data collection itself. This led to the development of MASSIF-1 at the ESRF, the world’s first beamline to run fully automatically while making intelligent decisions taking user requirements into account. Since opening in late 2014 the beamline has now processed over 39,000 samples. Improvements have been made to the speed of the sample handling robotics and error management within the software routines. The workflows initially put in place, while highly innovative at the time, have been expanded to include increased complexity and additional intelligence using the information gathered during characterisation, this includes adapting the beam diameter dynamically to match the diffraction volume within the crystal. Complex multi-position and multi-crystal data collections are now also integrated into the selection of experiments available. This has led to increased data quality and throughput allowing even the most challenging samples to be treated automatically.

**Synopsis**

Significant improvements in the sample location, characterisation and data collection algorithms on the autonomous ESRF beamline MASSIF-1 are described. The workflows now include dynamic beam diameter adjustment and multi-position and multi-crystal data collections.

## 1.0 Introduction

Automation is transforming the way scientific data are collected, allowing large amounts of high quality data to be gathered in a consistent manner (Quintana & Plätzer, 2015; Foster, 2005). Advances in robotics and software have been key in these developments and have had a particular impact on structural biology, allowing multiple constructs to be screened and purified (Camper & Viola, 2009; Hart & Waldo, 2013; Vijayachandran, *et al.*,2011); huge numbers of crystallisation experiments to be performed (Elsliger, *et al.*,2010; Ferrer *et.*, 2013; Heinemann, *et al.*, 2003; Joachimiak, 2009; Calero *et al.*, 2014), samples to be mounted at synchrotrons (Cipriani *et al.*, 2006; Cohen *et al.*, 2002; Jacquamet, *et al.*, 2009; Nurizzo, *et al.*, 2016; Papp, *et al.*, 2017; Snell, *et al.*, 2004), data to be analysed and processed (Bourenkov & Popov, 2010; Holton & Alber, 2004; Incardona, *et al.*, 2009; Leslie, *et al.*, 2002; Monaco, *et al.*, 2013; Winter, 2010) and the entire PDB to be validated (Joosten ei al., 2012). The combination of robotic sample mounting and on-line data analysis has been particularly important in macromolecular crystallography (MX) as it allowed time to be saved, large numbers of samples to be screened and enabled the remote operation of beamlines. However, despite these advances, a human presence is still required to sequence actions. Pioneering beamlines that fully automated the process, such as LRL-CAT at the APS (Wasserman, *et al.*, 2012) and the SSRL MX beamlines (Tsai, *et al.*, 2013), removed the need for human presence, but as they rely on optical loop centring this means that restrictions have to be placed on the size of crystals and tend to be for robust, well diffracting samples, generally those for proprietary research for the pharmaceutical industry.

In 2014 the ESRF beamline MASSIF-1(Bowler, *et al.*, 2015) opened to users as the first beamline to fully automate MX data collection, including sample location and complex decision making algorithms (Svensson, *et al.*, 2015). The combination of sample location and characterisation allows even the smallest and weakly diffracting samples to be treated automatically. This opened full automation to any sample presented in any mount and has provided a new tool to structural biologists, allowing the process of collecting hundreds of data sets or screening hundreds of crystals to be ‘outsourced’, freeing their time and often collecting better data (Bowler, *et al.*, 2016). At the time of writing, the beamline has processed more than 39,000 samples representing a wide range of projects, from those that require extensive screening to find the best diffracting crystal (Na, *et al.*, 2017; Sorigué *et al.*, 2017; Naschberger *et al.* 2017) to small molecule fragment screening (Cheeseman, *et al.*, 2017; Hiruma, *et al.*, 2017) and experimental phasing at high and low resolutions (Kharde, *et al.*, 2015; Muir, *et al.*, 2016). The beamline is able to deal with a wide range of samples by combining parameters provided by the user with information gathered during processing. The workflows initially put in place have performed very well but many enhancements remained possible.

Here, we describe how the algorithms have been improved to increase the amount and quality of data collected on MASSIF-1. Additional subroutines have been added to monitor and correct errors in centring and account for low resolution data collection as well as dynamically adjusting the beam diameter to match homogenous diffraction volumes. In combination with new multiple position and multiple crystal data collection workflows, fully automatic data collection is now possible for the most challenging samples.

## 2.0 Experimental details and results

### 2.1 Hardware improvements

One of the most time consuming, and important, steps in the X-ray centring process is the initial mesh scan that locates and characterises the crystal. When first implemented on MASSIF-1, a rotation of omega was included in the scan. This implementation required that triggering of the acquisition of images was instigated by the omega axis, meaning that each line of the mesh was treated as a separate data collection. The preparation required between data collections led to additional time being taken for the scan. We have now implemented a scan that includes no rotation of the omega axis and requires only the movements of the high precision Y/Z stage beneath the RoboDiff (Nurizzo, *et al.*, 2016). This allows the triggering of acquisition to be made by the motor position, allowing the whole mesh to be launched as a single data collection. This method was implemented in 2017 and comparison of the elapsed times for mesh scans performed in the last 2 months of 2016 with the first 2 months of 2017 shows that the time required for these scans has been reduced by an average of 1 minute (Figure 1).

**Figure 1.**
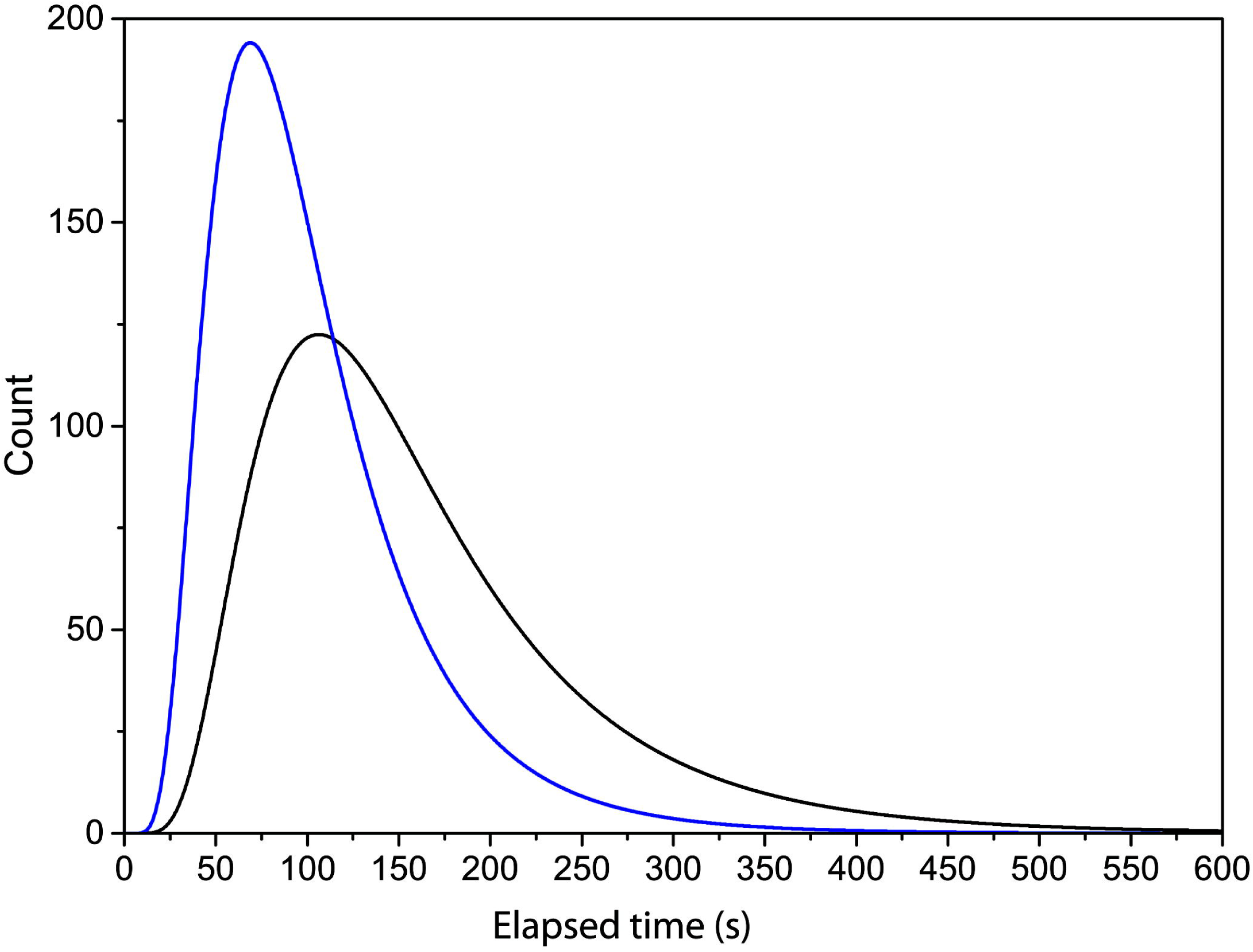
Decrease in time taken to perform the mesh scan after hardware improvements. Lognormal distribution of the elapsed time for mesh scans using the former protocol (black) and the new fast mesh (blue) for the three months preceding and following the new protocol.

### 2.2 Dynamic beam sizing

One of the benefits of running a completely automated system is the ability to collect large amounts of data on samples and use these data in improving strategies for data collection. We initially realised that the volumes of all crystals were determined during the X-ray centring routine and this information was subsequently included in the strategy calculation, having the biggest effect on the calculation of the maximum dose given to a crystal during data collection (Bowler, *et al.*, 2016; Svensson, *et al.*, 2015). Additionally, these measurements provided a distribution of crystal volumes allowing us to use a default beam diameter of 50 μm, as this was the crystal dimension most frequently observed on the beamline. A specific beam diameter can be selected on a per sample basis in the diffraction plan in ISPyB (Delagenière, *et al.*, 2011), however, this option is usually used when users are sure that crystal volumes are significantly smaller than the default beam diameter (Figure 2). Using the information gathered during the mesh scan we can determine an optimised beam diameter. By accurately matching beam diameter to the crystal, it has been shown that the background can be dramatically reduced (Holton & Frankel, 2010; Moukhametzianov, *et al.*, 2008). This is particularly striking when crystals are very small (Evans, *et al.*, 2011) but if a crystal is large, the additional diffraction power should not be wasted. This also has to be balanced with the degree of variability within each crystal (Bowler & Bowler, 2014; Bowler, *et al.*, 2010; Pozharski, 2012).

**Figure 2.**
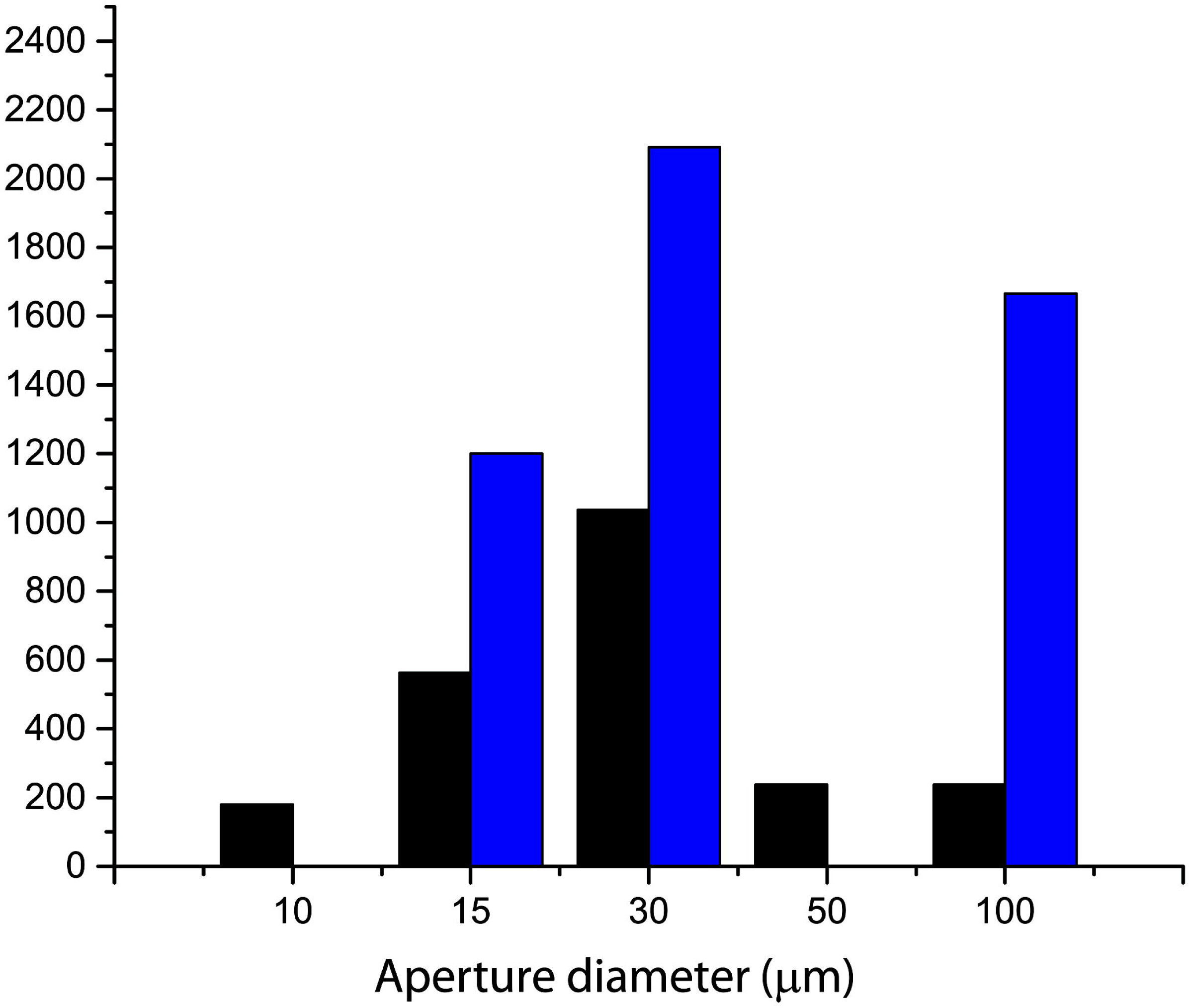
Beam diameter selections by users and algorithm in 2017. The number of times a beam diameter is selected either by the user (black) or automatically (blue) is shown. 6877 data collections were performed with a beam diameter at the default value of 50 μm, as this is not changed by the software, the value is not shown.

We have now introduced a dynamic beam diameter adjustment into all workflows running on MASSIF-1 where no value has been pre-selected by the user. The X-ray centring routine determines the crystal position relative to the beam, the crystal dimensions and also determines the best homogenous volume within the crystal. The centre of mass of this volume is then used as the centring position and it is the dimensions of this volume that are used to select the beam diameter. There are 5 beam diameters available on MASSIF-1, 100 μm, 50 μm, 30 μm, 15 μm and 10 μm (Bowler, *et al.*, 2015) and the smallest vertical volume dimension is used to select the aperture that matches most closely. In this way, the largest volume can be illuminated without increasing background or ‘contaminating’ the diffraction from variable areas. All steps during X-ray centring are performed with the 50 μm aperture. As the scans are performed with an overlap, the smallest dimension that can be measured is ~20 μm meaning that the 10 μm option is only used when selected by the user. Once X-ray centring is completed, the new aperture diameter is selected and characterisation images are collected using this diameter. The strategy calculation will include the new flux for the aperture as well as the crystal volume determined during the centring procedure. Since introducing the adaptable beam diameter the system has selected the 30μm beam size most frequently (Figure 2) followed by the 100 μm and the 15 μm diameter. Half of all data collections are performed with a diameter of 50 μm, reinforcing its choice as the default value.

Can the advantages of dynamic beam adjustment be demonstrated in a consistent manner? It is always problematic to clearly show that one data collection method is better than another. However, here we show that dynamic beam sizing make a significant difference for weakly diffracting samples.

We initially tested the adaptive beam diameter protocol on crystals of the β_1_-adregenic GPCR (Warne, *et al.*, 2008). These crystals diffract weakly, exhibit considerable variation in diffraction quality and tend to form as thin plates or needles. A total of 30 crystals were run on MASSIF-1, first using the classic protocol (Svensson, *et al.*, 2015), where a 50 μm beam diameter is the default, and then running a second protocol on the same crystal including the adaptive beam diameter. In many cases, data sets were collected from the same crystal using both procedures; however, for some cases, data sets were only processed where the beam diameter had been reduced to match the crystal size. Table 1 shows crystal dimensions and data processing statistics from crystals where an automatically processed data set was produced (8 out of 30 crystals). Where crystals were of sufficient quality, data sets were mostly produced from both protocols, but it is for the smaller crystals that a difference is discernible. For crystals with a *y* dimension below 30 μm the data sets produced have a higher <I/σ(I)> or resolution limit (Table 1; For42, For48, For59 and For67) even though the crystals have already been exposed. For one of these crystals, For42, the data set collected with the smaller beam is significantly better (Table 2).

**Table 1.** **Data collection details for β_1_-andregenic GPCR crystal with standard and adapted beam diameter protocols.** The dimensions are: *x* the measured crystal length paralell to the spindle axis, *y* the height orthogonal to the spindle axis and *z* the depth orthogonal to the spindle axis 90° away in ω.

| Crystal | Crystal dimensions ($x$, $y$, $z$, mm) | Fixed beam diameter | | Adaptable beam diameter | |
| --- | --- | --- | --- | --- | --- |
| | | Resolution limit ($\text{\AA}$) | $\langle I/\sigma(I) \rangle$ | Resolution limit ($\text{\AA}$) | $\langle I/\sigma(I) \rangle$ |
| adrcpt-For41 | 0.109 x 0.053 x 0.025 | 3.77 | 6.7 | 4.13 | 4.4 |
| adrcpt-For42 | 0.084 x 0.025 x 0.025 | 4.22 | 4.3 | 3.53 | 10.6 |
| adrcpt-For45 | 0.035 x 0.045 x 0.051 | 3.95 | 6.2 | - | - |
| adrcpt-For47 | 0.105 x 0.061 x 0.051 | 3.74 | 4.7 | 3.72 | 5.4 |
| adrcpt-For48 | 0.105 x 0.039 x 0.064 | - | - | 3.8 | 5.7 |
| adrcpt-For58 | 0.169 x 0.050 x 0.061 | 3.88 | 6.6 | 4.11 | 4.5 |
| adrcpt-For59 | 0.042 x 0.024 x 0.025 | 3.25 | 9.2 | 3.16 | 8.3 |
| adrcpt-For67 | 0.064 x 0.026 x 0.031 | - | - | 3.8 | 5.6 |

**Table 2.** **Comparison of data collection strategies and processing statistics from standard and adaptive beam diamter protocols for a β_1_-andregenic GPCR crystal.**

| Crystal | adrcpt-For42 | adrcpt-For42 |
| --- | --- | --- |
| Beam diameter ($\mu\text{m}$) | 50 | 30 |
| Space group (no.) | $P2_12_12_1$ (19) | $P2_12_12_1$ (19) |
| Cell parameters ($\text{\AA}$, $^\circ$) | 116.8, 121.1, 129.5<br>90, 90, 90 | 116.5, 120.78, 128.7<br>90, 90, 90 |
| Flux (ph/s) | $5.7 \times 10^{11}$ | $2.2 \times 10^{11}$ |
| Rotation width ($^\circ$) | 0.15 | 0.05 |
| Total oscillation Range ($^\circ$) | 149.1 | 124.0 |
| Total dose (MGy) | 5.25 | 5.92 |
| Detector resolution | 4.05 | 3.9 |
| Wilson B-factor ($\text{\AA}^2$) | 94.0 | 97.0 |
| Resolution ($\text{\AA}$) | 48.2 – 3.95 (4.09 – 3.95) | 48.7 – 3.53 (3.66-3.53) |
| Completeness (%) | 98.2 (90.9) | 81.5 (33.0) |
| Observed reflections | 15,940 (1413) | 18,785 (721) |
| Average redundancy | 4.1 (2.7) | 4.4 (1.6) |
| $\langle I/\sigma(I) \rangle$ | 4.3 (0.8) | 10.6 (0.6) |
| $R_{\text{meas}}$ | 0.23 (1.45) | 0.1 (1.64) |
| $R_{\text{merge}}$ | 0.18 (1.05) | 0.077 (1.17) |
| $CC_{1/2}$ | 0.99 (0.43) | 1 (0.33) |

What effect has the protocol had on overall data collection? In order to analyse the difference we looked at the average signal to noise ratios, <I/σ(I)>, for all data sets processed automatically (Monaco, *et al.*, 2013; Vonrhein, *et al.*, 2011; Sparta, *et al.*, 2016; Winter, 2010) for the year preceding and following the introduction of the protocol. This amounts to data for approximately 22,000 samples. Figure 3 shows the distributions of overall <I/σ(I)>for the data sets. While the distributions are similar for high <I/σ(I)> they diverge at the lower values with a significant shift lower for the adaptive beam dimeter data sets. The average before the procedure was put in place is 14.4, moving down to 12.2 after, with the modal values 8.55 before and 5.9 after (Figure 3.). We initially found this surprising as we had expected a general increase in <I/σ(I)>. However, given the effect seen on the GPCR crystal, the distribution change is understandable. While the beam diameter is increased or decreased to match the diffraction volume, the <I/σ(I)> values for strongly diffracting crystals will remain the same given the dose to achieve a certain resolution without radiation damage, taking the crystal volume and changed flux into account. However, it is for the weakly diffracting crystals that the adaptive beam dimeter has the most significant effect. Whether the diameter is increased or decreased there is a large shift in the number of data sets that are processed that have rather low <I/σ(I)> after the adaptive beam diameter protocol was implemented. This implies that by introducing this routine into the regular data collection workflow, the beamline is able to increase the number of data sets processed from these samples by reducing the background noise.

**Figure 3.**
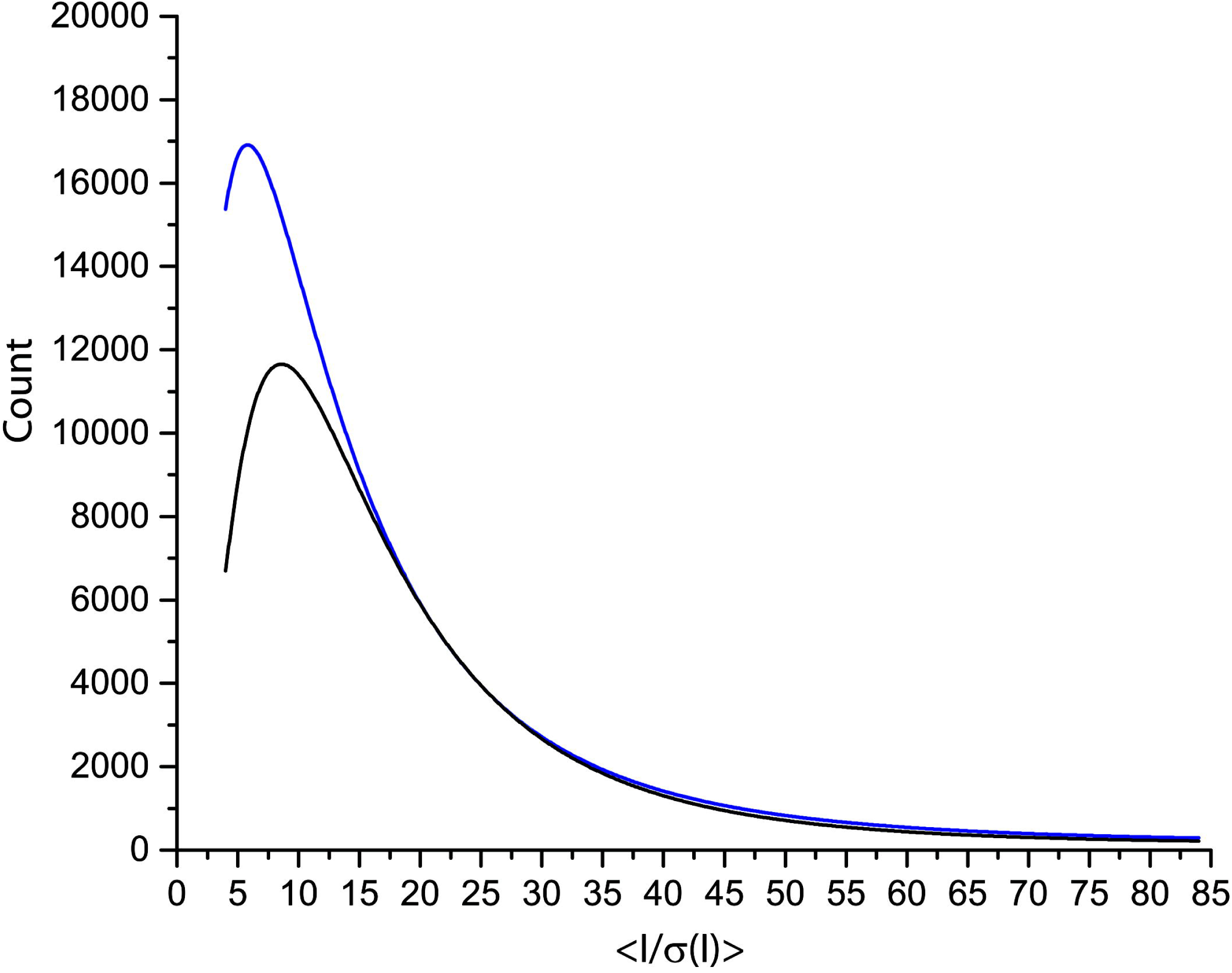
Distribution of overall <I/σ(I)> for data sets processed from the year preceding dynamic beam aperture (black) and the year after dynamic beam aperture (blue). There is a significant shift in the number of data sets processed with lower <I/σ(I)> after dynamic beam adjustment was introduced.

### 2.3 Improved error handling

The correct handling of errors is paramount in an automated system. We initially introduced many error handling routines at both a high level, such as the collection of a data set with default settings when indexing fails (Svensson, *et al.*, 2015), and a low level, such as escaping from small robotic errors (Nurizzo, *et al.*, 2016). After processing more than 39,000 samples we have now been able to observe most errors encountered and have extended the processes to catch them.

#### 2.3.1 Centring errors

We have occasionally observed that after the X-ray centring routine the crystal was still not correctly aligned over the full rotation range. This may be due to movements of the support after the routine is completed or to errors in the routine arising from multiple peaks being selected for centring. Whatever the reason, it can lead to a data set being lost. We therefore introduced a check in the characterisation step that ensures that the 4 images have a diffraction signal. If one image has a signal that is below 10% of the top signal, a recovery routine is launched. This involves three short line scans (50 μm above and below the current centred position) being launched over the currently centred position. In most cases it corrects the error. In 2017, 13,776 samples were processed on MASSIF-1 and centring recovery was launched 221 times. This represents a centring error rate of 1.6% which should be reduced further by being able to detect and recover incorrectly centred samples.

#### 2.3.2 Low resolution data collection

Unless a resolution is specified by the user, all mesh and characterisation images are collected at 2 Å. If the predicted resolution extends beyond the corners of the detector (1.42 Å) the detector is moved to the new resolution and a further characterisation is launched. This allows the highest possible resolution to be obtained and ensures that characterisation is performed at an optimal detector distance (Svensson, *et al.*, 2015). However, for low resolution, the resolution determined by BEST is used and data collection proceeds according to the determined strategy. It would seem sensible that if very low diffraction is determined the characterisation images should also be re-collected at this resolution. We therefore introduced a routine to re-collect the characterisation images at 4 Å for all samples where the determined resolution is below this value. This allows the distribution of intensity to be better estimated and should lead to better strategy calculations (Popov & Bourenkov, 2003).

By analysing the relationship between predicted and determined resolution from all data sets collected so far we can also try to improve the quality of data collected on MASSIF-1 (Figure 4). The distribution shows that the agreement is excellent, usually slightly underestimating the achievable resolution. This may well be due to the difference between criterion for resolution limit determination being <I/σ(I)> for characterisation and CC1/2 for complete data sets. A clear trend is that for weakly diffracting crystals the strategy tends to underestimate the resolution (Figure 4). This is due to the difficulty in estimating a B factor at very low resolution. In addition to re-collecting the characterisation images the procedure now always sets the detector resolution to 4 Å for all data collections where the predicted resolution is lower than this value. In this way, we hope that higher resolution data will not be missed, if possible, as complete data to 4 Å is more important than sub-optimal data collection at 7 Å.

**Figure 4.**
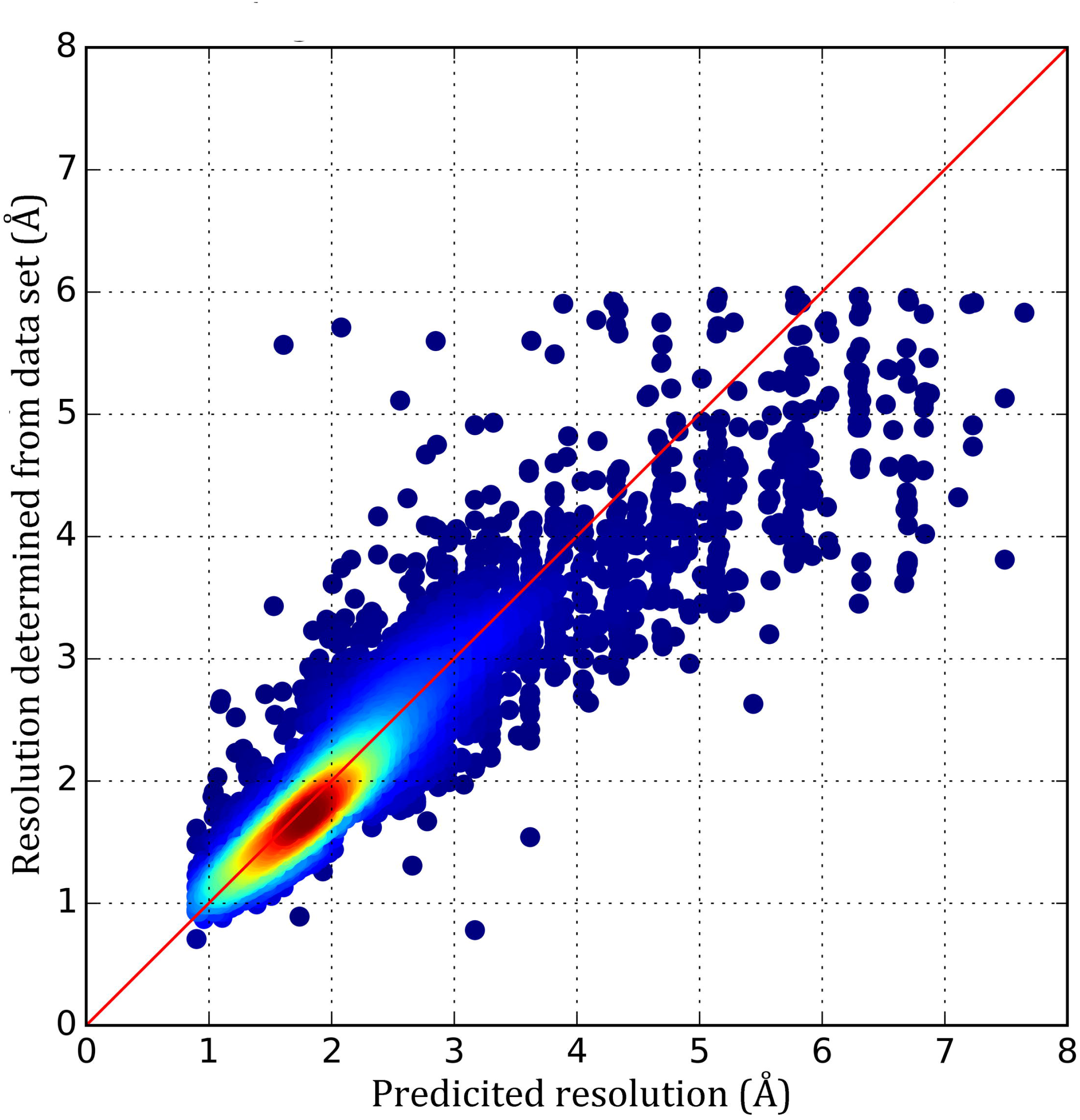
Scatter plot of predicied resolutions for data collections against the resolution determined by autoprocessing for all crystals processed so far on MASSIF-1 that resulted in an automatically processed data set. The red line shows perfect agreement between predicted and achieved resolution and the gradient shows the density of data points.

### 2.4 Multiple crystal and multiple position data collection strategies

The possibility to input a number of positions from which to collect data was introduced into the diffraction plan early in the operation of MASSIF-1 (Bowler, *et al.*, 2016; Svensson *et al.*, 2015). This has allowed complete data sets to be collected either from separate crystals contained on a sample support or from multiple positions within a single crystal and has proved to be a popular option with many samples received on MASSIF-1 having a number of positions between 2 and 12 requested (Figure 5). While extremely useful, this protocol does not cover the scenario where radiation sensitive samples can benefit from a large dose being spread over multiple partial data sets, a procedure known generally as helical data collection (Flot *et al,* 2010) that has been shown to be beneficial in many cases (Polsinelli, *et al.*, 2017). Radiation damage can often make it difficult to collect complete data, or data with sufficient anomalous signal, from a single crystal or a single position within a crystal. A new experiment type is now available that will automatically collect multiple partial data sets from positions within a homogenous volume of a crystal. This can lead to improved data quality, increased resolution and higher anomalous peaks. This is the first fully automated helical data collection protocol that also accounts for the heterogeneity of crystal diffraction quality.

**Figure 5.**
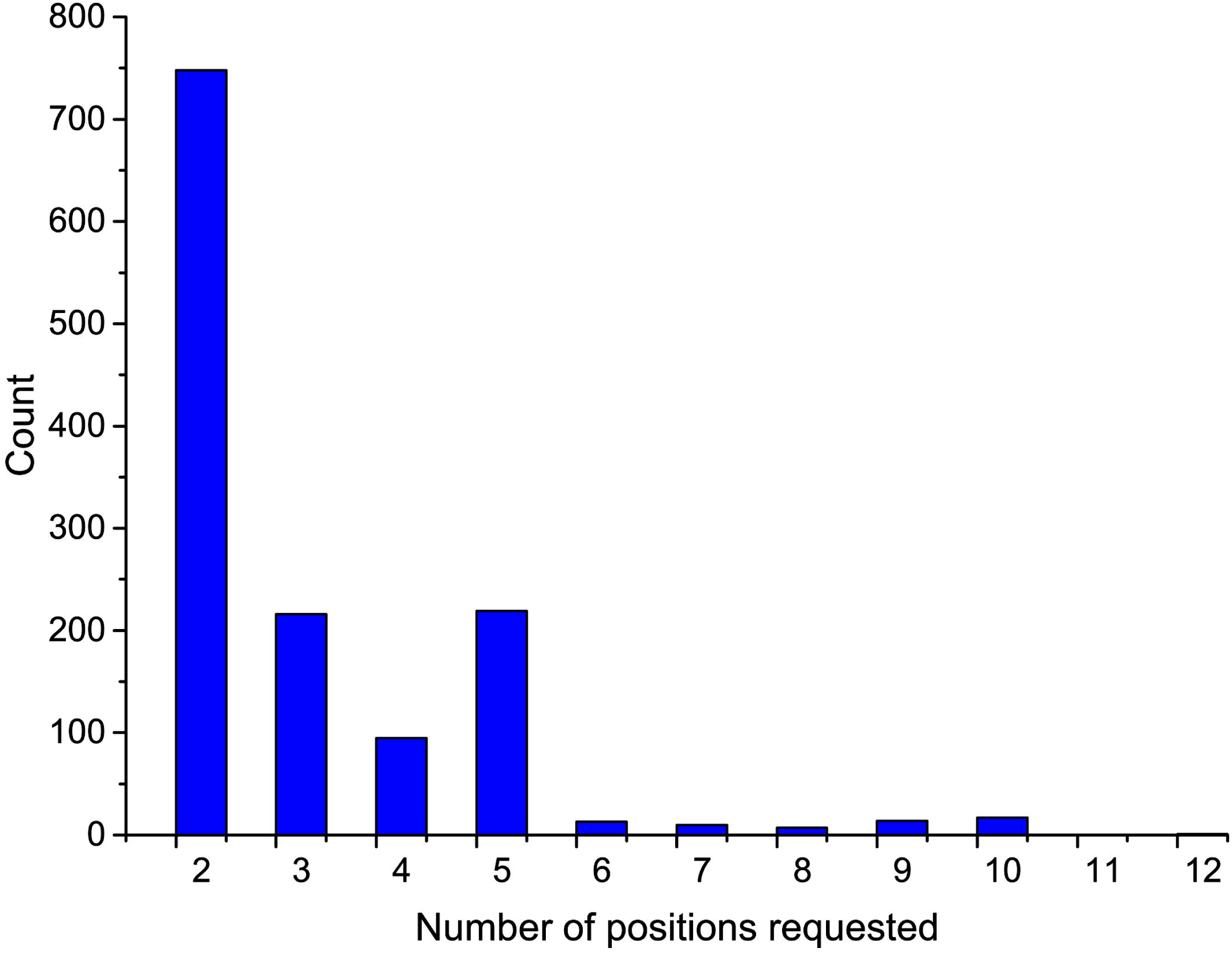
Number of positions selected by users for multiple crystal and multiple position data collections in 2017. Multiple position data collections are requested for 9% of samples for this year.

**Figure 6.**
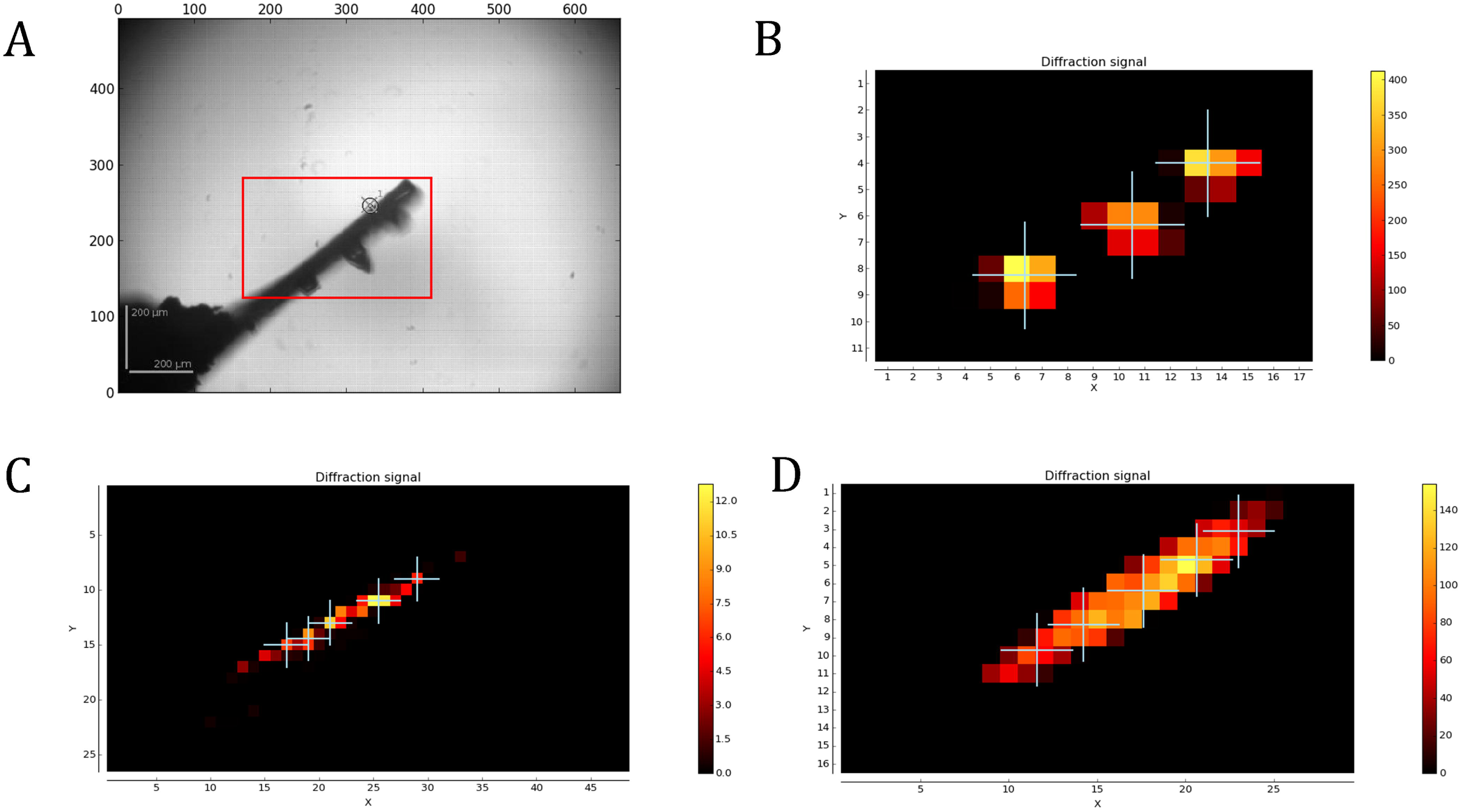
Multiple crystal and position data collection. **A.** Automesh of a CrystalDirect (Zander, Hoffmann, *et al.*, 2016) support that has 3 crystals mounted. The widest orientation of the mount was selected. **B**. Mesh scan of the mount shown in A. Three positions were requested and three detected. **C**. Mesh scan for a βPGM crystal where a native pseudo-helical data collection was requested, 5 positions were detected and a beam diamter of 30 μm selected and **D** Mesh scan for a FAE crystal where a SAD pseudo-helical data collection was requested, 5 positions were detected and a beam diamter of 100 μm selected.

Multiple position/crystal experiments are selected either by specifying a number of positions in the diffraction plan for the sample or by requesting MXPressP or MXPressP_SAD (for *Pseudo helical* and *Pseudo helical with SAD strategy*) in the ‘experiment type’ field for the required samples in ISPyB (Delagenière *et al.*, 2011). These new experiments operate in much the same way as the usual automated workflows on MASSIF-1 in that all the current features are retained, such as resolution selection, strategy input, diffraction volume calculation and smart beam sizing. If multiple positions are selected, the *automesh* algorithm, that determines the area to scan to locate the crystal (Svensson *et al.*, 2015), uses the widest orientation of the sample support, rather than the smallest, in order to avoid overlapping crystals or positions in ω. After the mesh scan is complete, the map is analysed either for the number of peaks requested (if no positions are specified for MXPressP the default is 5). Determined peaks must be within 10% of the value of the peak for multiple crystals or 70% of the second highest peak for pseudo helical. Additionally, any peaks that are closer than a beam diameter together or that will overlap in ω are eliminated. The number of allowed detected peaks is then specified by a comment in ISPyB. If multiple crystals have been selected, each point is then centred as usual and a complete data set collected according to user input requirements. If MXPressP is selected, the top peak is centred and 4 characterisation images are collected from the best position. A strategy is then calculated for a complete data set and the data collected, for MXPressP_SAD the strategy is optimised for structure solution by SAD (Svensson *et al.*, 2015). As usual, in case of indexing failure, a default data collection of 180° is collected (240° for triclinic and 360° for SAD data collection). Once completed, a strategy is then calculated to collect a complete data set from the *N* positions determined in the mesh scan that are within 70% of the value of position 2. The strategy, as usual, accounts for the volume of the positions, beam diameter etc. Again, in case of a failure in indexing, default data collections are performed at each position using a full dose and the rotation range determined by 180°/*N* (240°/*N* for triclinic and 360°/*N* for SAD). Each partial data set has a 5° overlap with the next to assist with scaling.

We are, for the moment, remaining cautions with helical data collection by collecting a full single position complete data set from the best volume. The reason for this is twofold: 1. we have observed that crystal heterogeneity can often lead to a number of the partial data sets being of varying quality despite the stringent quality threshold we have implemented and 2. We are eager to compile a large amount of data on how and when helical data collection is superior to single position. This is extremely important as, so far, the few studies on helical data collection have not considered crystal heterogeneity (Bowler & Bowler, 2014; Bowler *et al.*, 2010). Strategy parameters and data processing statistics for two example systems using the pseudo helical routines for native and SAD data collections are shown in Table 3. Two proteins that tend to form crystals with a needle morphology were selected: β-phosphoglucomutase (βPGM) in an open conformation (Baxter *et al.*, 2010) and ferulic acid esterase (FAE) that contains 8 × Se and 5 × Cd^2+^ (Prates *et al.*, 2001) with a significant anomalous signal at the MASSIF-1 wavelength of 0.966 Å. Comparing the single position data collection to the merged multiple position data sets shows that in these cases there is not a significant increase in data quality. However, in the SAD case the helical data set has considerably higher <I/σ(I)>, anomalous correlation coefficients and mid-slope of anomalous probability than the single position data set. For the native data sets, the single position is slightly better. This may reflect the heterogeneity within the crystal and highlights the importance of this parameter in whether to select helical versus single position for a certain project. The ability to automatically run clustering algorithms (Giordano *et al.*, 2012; Zander, Cianci, *et al.*, 2016) on these partial data sets may also improve the quality of the final data. We hope that by being able to analyse the variation in diffraction quality, and compare single position data to multi-position data from the same crystal, a more general strategy for these types of data collection may emerge.

**Table 3.**
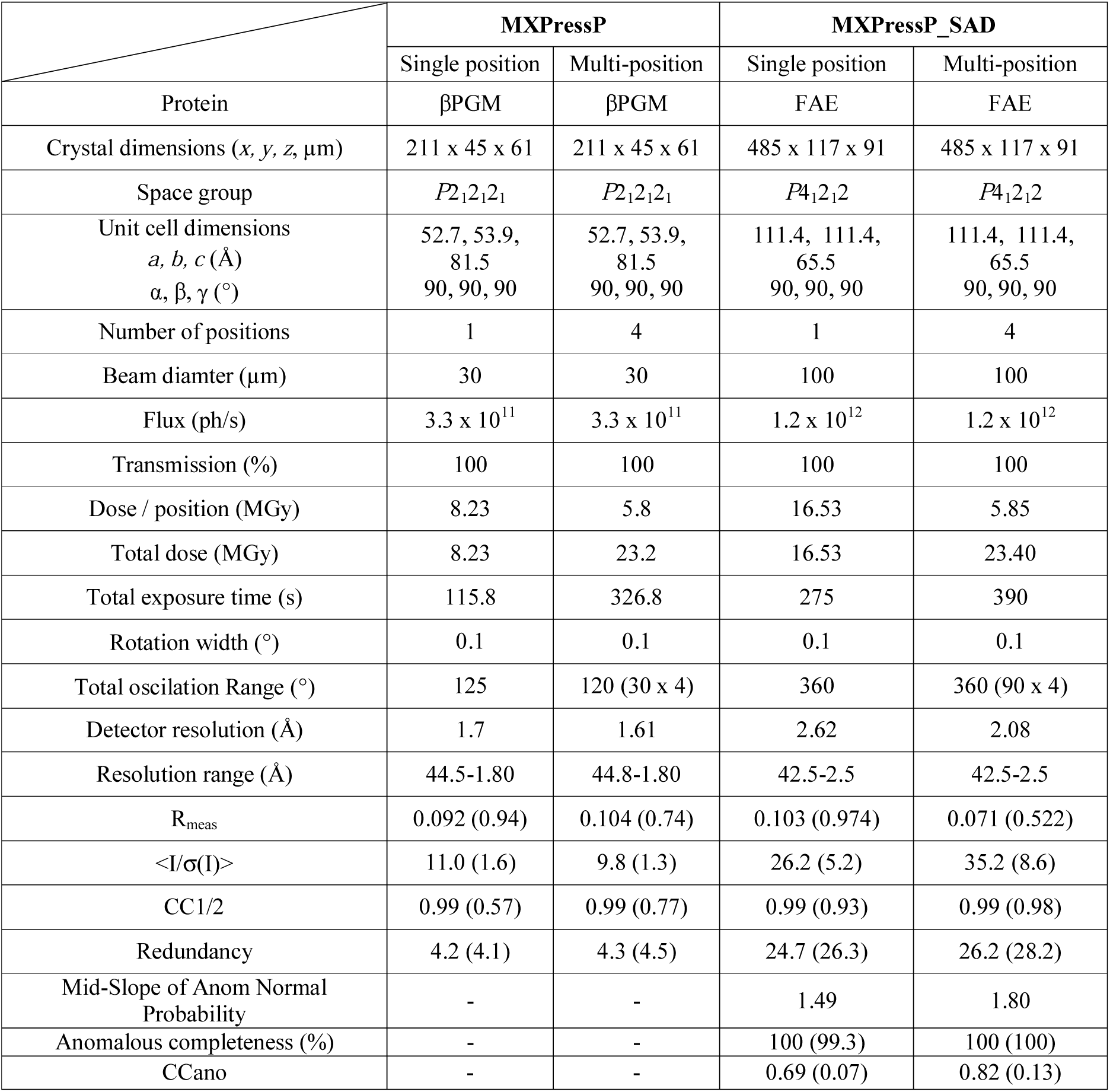
**Pseudo helical data collection.** Comparison of data sets collected from a single position and from multiple positions for native and SAD strategies.

## 3.0 Discussion

The results presented here demonstrate not only the increase in the speed and reliability of automatic data collections but also that more complex strategies can be brought into the arena of autonomous experiments. Automation is often seen as a way to deal with mundane experiments that require little human input. The autonomous system presented here is different in that, in addition to automating mounting and centring, it also uses data gathered during the process to improve data collection strategies. We have already demonstrated that MASSIF-1 collects, on average, better quality data than humans are able to (Bowler *et al.*, 2016). The additional routines presented here add even more intelligence into the system that should further enhance its ability to extract the best possible data from every sample. This built-in intelligence means that the system is excellent for not only robust and routine data collections but also for challenging systems that diffract weakly. We have demonstrated that adapting the beam dimeter can increase the number of data sets that can be processed from these types of sample. We hope that by providing more data on more samples we can improve feedback into experiment cycles and increase the amount of useful data produced.

All the developments described here have been exported to the human operated ESRF beamlines (Mueller-Dieckmann *et al.*, 2015). As structural biologists now turn to a an ever wider variety of techniques, we hope that fully automatic data collection will become the standard data collection method for MX as the best possible data can be collected from samples, be they large and robust or small and weakly diffracting. In combination with developments in the robotic mounting and soaking of crystals (Zander, Hoffmann, *et al.*, 2016) we envision that the future of macromolecular crystallography is the provision of a fully automated high throughput service able to rapidly produce high quality structural models and screen for potential therapeutic and probe molecules.

## Acknowledgments

We thank Tony Warne (MRC-LMB, Cambridge, UK) for the gift of β1-androgenic GPCR crystals; Josan Marquez and Guillaume Hoffman (EMBL, Grenoble) for providing CrystalDirect supports with multiple crystals; and the ESRF Automation Task Force (ATF) and the ESRF-EMBL Joint Structural Biology Group (JSBG) for supporting the developments. We are also grateful to Gordon Leonard (ESRF) for comments on the manuscript.

## References

Baxter, N. J., Bowler, M. W., Alizadeh, T., Cliff, M. J., Hounslow, A. M., Wu, B., Berkowitz, D. B., Williams, N. H., Blackburn, G. M. & Waltho, J. P. (2010). Atomic details of near-transition state conformers for enzyme phosphoryl transfer revealed by MgF-3 rather than by phosphoranes, Proc Natl Acad Sci U S A 107, 4555–4560.

Bourenkov, G. P. & Popov, A. N. (2010). Optimization of data collection taking radiation damage into account, Acta Cryst. D 66, 409–419.

Bowler, M.W., Svensson, O. & Nurizzo, D. (2016). Fully automatic macromolecular crystallography: the impact of MASSIF-1 on the optimum acquisition and quality of data, Cryst. Rev., 22, 233–249.

Bowler, M. G. & Bowler, M. W. (2014). Measurement of the intrinsic variability within protein crystals: implications for sample-evaluation and data-collection strategies, Acta Cryst. F 70, 127–132.

Bowler, M. W., Guijarro, M., Petitdemange, S., Baker, I., Svensson, O., Burghammer, M., Mueller-Dieckmann, C., Gordon, E. J., Flot, D., McSweeney, S. M. & Leonard, G. A. (2010). Diffraction cartography: applying microbeams to macromolecular crystallography sample evaluation and data collection, Acta Cryst. D 66, 855–864.

Bowler, M. W., Nurizzo, D., Barrett, R., Beteva, A., Bodin, M., Caserotto, H., Delageniere, S., Dobias, F., Flot, D., Giraud, T., Guichard, N., Guijarro, M., Lentini, M., Leonard, G. A., McSweeney, S., Oskarsson, M., Schmidt, W., Snigirev, A., von Stetten, D., Surr, J., Svensson, O., Theveneau, P. & Mueller-Dieckmann, C. (2015). MASSIF-1: a beamline dedicated to the fully automatic characterization and data collection from crystals of biological macromolecules, J Synch. Rad. 22, 1540–1547.

Calero, G., Cohen, A. E., Luft, J. R., Newman, J. & Snell, E. H. (2014). Identifying, studying and making good use of macromolecular crystals, Acta Cryst. F70, 993–1008.

Camper, D. V. & Viola, R. E. (2009). Fully automated protein purification, Anal. Biochem. 393, 176–181.

Cheeseman, M. D., Chessum, N. E. A., Rye, C. S., Pasqua, A. E., Tucker, M. J., Wilding, B., Evans, L. E., Lepri, S., Richards, M., Sharp, S. Y., Ali, S., Rowlands, M., O’Fee, L., Miah, A., Hayes, A., Henley, A. T., Powers, M., te Poele, R., De Billy, E., Pellegrino, L., Raynaud, F., Burke, R., van Montfort, R. L. M., Eccles, S. A., Workman, P. & Jones, K. (2017). Discovery of a Chemical Probe Bisamide (CCT251236): An Orally Bioavailable Efficacious Pirin Ligand from a Heat Shock Transcription Factor 1 (HSF1) Phenotypic Screen, J. Med. Chem. 60, 180–201.

Cipriani, F., Felisaz, F., Launer, L., Aksoy, J. S., Caserotto, H., Cusack, S., Dallery, M., di-Chiaro, F., Guijarro, M., Huet, J., Larsen, S., Lentini, M., McCarthy, J., McSweeney, S., Ravelli, R., Renier, M., Taffut, C., Thompson, A., Leonard, G. A. & Walsh, M. A. (2006). Automation of sample mounting for macromolecular crystallography, Acta Cryst. D62, 1251–1259.

Cohen, A. E., Ellis, P. J., Miller, M. D., Deacon, A. M. & Phizackerley, R. P. (2002). An automated system to mount cryo-cooled protein crystals on a synchrotron beamline, using compact sample cassettes and a small-scale robot, J. App. Cryst. 35, 720–726.

Delagenière, S., Brenchereau, P., Launer, L., Ashton, A. W., Leal, R., Veyrier, S., Gabadinho, J., Gordon, E. J., Jones, S. D., Levik, K. E., McSweeney, S. M., Monaco, S., Nanao, M., Spruce, D., Svensson, O., Walsh, M. A. & Leonard, G. A. (2011). ISPyB: an Information Management System for Synchrotron Macromolecular Crystallography, Bioinformatics 27, 3186–3192.

Elsliger, M.-A., Deacon, A. M., Godzik, A., Lesley, S. A., Wooley, J., Wuthrich, K. & Wilson, I. A. (2010). The JCSG high-throughput structural biology pipeline, Acta Cryst. F66, 1137–1142.

Evans, G., Axford, D. & Owen, R. L. (2011). The design of macromolecular crystallography diffraction experiments, Acta Cryst. D 67, 261–270.

Ferrer, J. L., Larive, N. A., Bowler, M. W. & Nurizzo, D. (2013). Recent progress in robot-based systems for crystallography and their contribution to drug discovery, Exp. Opin. Drug;. Disc. 8, 835–847.

Flot, D., Mairs, T., Giraud, T., Guijarro, M., Lesourd, M., Rey, V., van Brussel, D., Morawe, C., Borel, C., Hignette, O., Chavanne, J., Nurizzo, D., McSweeney, S. & Mitchell, E. (2010). The ID23-2 structural biology microfocus beamline at the ESRF, J. Synch. Rad. 17, 107–118.

Foster, I. (2005). Service-Oriented Science, Science 308, 814–817.

Giordano, R., Leal, R. M. F., Bourenkov, G. P., McSweeney, S. & Popov, A. N. (2012). The application of hierarchical cluster analysis to the selection of isomorphous crystals, Acta Cryst. D 68, 649–658.

Hart, D. J. & Waldo, G. S. (2013). Library methods for structural biology of challenging proteins and their complexes, Curr.Opin. Struct. Biol. 23, 403–408.

Heinemann, U., Bussow, K., Mueller, U. & Umbach, P. (2003). Facilities and methods for the high-throughput crystal structural analysis of human proteins, Acc. Chem. Res. 36, 157–163.

Hiruma, Y., Koch, A., Hazraty, N., Tsakou, F., Medema, R. H., Joosten, R. P. & Perrakis, A. (2017). Understanding inhibitor resistance in Mps1 kinase through novel biophysical assays and structures, J. Biol. Chem. 292, 14496–14504

Holton, J. & Alber, T. (2004). Automated protein crystal structure determination using elves, Proc. Natl. Acad. Sci. USA 101, 1537–1542.

Holton, J. M. & Frankel, K. A. (2010). The minimum crystal size needed for a complete diffraction data set, Acta Cryst. D66, 393–408.

Incardona, M. F., Bourenkov, G. P., Levik, K., Pieritz, R. A., Popov, A. N. & Svensson, O. (2009). EDNA: a framework for plugin-based applications applied to X-ray experiment online data analysis, J. Synch. Rad. 16, 872–879.

Jacquamet, L., Joly, J., Bertoni, A., Charrault, P., Pirocchi, M., Vernede, X., Bouis, F., Borel, F., Perin, J. P., Denis, T., Rechatin, J. L. & Ferrer, J. L. (2009). Upgrade of the CATS sample changer on FIP-BM30A at the ESRF: towards a commercialized standard, J. Synch. Rad. 16, 14–21.

Joachimiak, A. (2009). High-throughput crystallography for structural genomics, Curr. Opin. Struct. Biol. 19, 573–584.

Joosten, R. P., Joosten, K., Murshudov, G. N. & Perrakis, A. (2012). PDB_REDO: constructive validation, more than just looking for errors, Acta Cryst. D 68, 484–496.

Kharde, S., Calviño, F. R., Gumiero, A., Wild, K. & Sinning, I. (2015). The structure of Rpf2-Rrs1 explains its role in ribosome biogenesis, Nucleic Acids Res. 43, 7083–7095.

Leslie, A. G., Powell, H. R., Winter, G., Svensson, O., Spruce, D., McSweeney, S., Love, D., Kinder, S., Duke, E. & Nave, C. (2002). Automation of the collection and processing of X-ray diffraction data -- a generic approach, Acta Cryst. D 58, 1924–1928.

Monaco, S., Gordon, E., Bowler, M. W., Delageniere, S., Guijarro, M., Spruce, D., Svensson, O., McSweeney, S. M., McCarthy, A. A., Leonard, G. & Nanao, M. H. (2013). Automatic processing of macromolecular crystallography X-ray diffraction data at the ESRF, J. App. Cryst. 46, 804–810.

Moukhametzianov, R., Burghammer, M., Edwards, P. C., Petitdemange, S., Popov, D., Fransen, M., McMullan, G., Schertler, G. F. X. & Riekel, C. (2008). Protein crystallography with a micrometre-sized synchrotron-radiation beam, Acta Cryst. D 64, 158–166.

Mueller-Dieckmann, C., Bowler, M., Carpentier, P., Flot, D., McCarthy, A., Nanao, M., Nurizzo, D., Pernot, P., Popov, A., Round, A., Royant, A., de Sanctis, D., von Stetten, D. & Leonard, G. (2015). The status of the macromolecular crystallography beamlines at the European Synchrotron Radiation Facility, Eur. Phys. J. Plus 130, 1–11.

Muir, Kyle W., Kschonsak, M., Li, Y., Metz, J., Haering, Christian H. & Panne, D. (2016). Structure of the Pds5-Scc1 Complex and Implications for Cohesin Function, Cell Rep. 14, 2116–2126.

Na, Z., Yeo, S. P., Bharath, S. R., Bowler, M. W., Balikci, E., Wang, C.-I. & Song, H. (2017). Structural basis for blocking PD-1-mediated immune suppression by therapeutic antibody pembrolizumab, Cell Res 27, 147–150.

Naschberger, A., Orry, A., Lechner, S., Bowler, M. W., Nurizzo, D., Novokmet, M., Keller, M. A., Oemer, G., Seppi, D., Haslbeck, M., Pansi, K., Dieplinger, H. & Rupp, B. (2017) Structural Evidence for a Role of the Multi-functional Human Glycoprotein Afamin in Wnt Transport, Structure 25, 1907–1915.

Nurizzo, D., Bowler, M. W., Caserotto, H., Dobias, F., Giraud, T., Surr, J., Guichard, N., Papp, G., Guijarro, M., Mueller-Dieckmann, C., Flot, D., McSweeney, S., Cipriani, F., Theveneau, P. & Leonard, G. A. (2016). RoboDiff: combining a sample changer and goniometer for highly automated macromolecular crystallography experiments, Acta Cryst. D 72, 966–975.

Papp, G., Felisaz, F., Sorez, C., Lopez-Marrero, M., Janocha, R., Manjasetty, B., Gobbo, A., Belrhali, H., Bowler, M. W. & Cipriani, F. (2017). FlexED8: the first member of a fast and flexible sample-changer family for macromolecular crystallography, Acta Cryst. D 73, 841–851.

Polsinelli, I., Savko, M., Rouanet-Mehouas, C., Ciccone, L., Nencetti, S., Orlandini, E., Stura, E. A. & Shepard, W. (2017). Comparison of helical scan and standard rotation methods in single-crystal X-ray data collection strategies, J. Synch. Rad. 24, 42–52.

Popov, A. N. & Bourenkov, G. P. (2003). Choice of data-collection parameters based on statistic modelling, Acta Cryst. D59, 1145–1153.

Pozharski, E. (2012). On the variability of experimental data in macromolecular crystallography, Acta Cryst. D68, 1077–1087.

Prates, J. A. M., Tarbouriech, N., Charnock, S. J., Fontes, C. M. G. A., Ferreira, L. s. M. A. & Davies, G. J. (2001). The Structure of the Feruloyl Esterase Module of Xylanase 10B from Clostridium thermocellum Provides Insights into Substrate Recognition, Structure 9, 1183–1190.

Quintana, F. J. & Plätzer, K. (2015). Addressing challenges in data collection: The role of automation in complex translational research, Science 349, 1567–1567.

Snell, G., Cork, C., Nordmeyer, R., Cornell, E., Meigs, G., Yegian, D., Jaklevic, J., Jin, J., Stevens, R. C. & Earnest, T. (2004). Automated sample mounting and alignment system for biological crystallography at a synchrotron source, Structure 12, 537–545.

Sorigué, D., Légeret, B., Cuiné, S., Blangy, S., Moulin, S., Billon, E., Richaud, P., Brugière, S., Couté, Y., Nurizzo, D., Müller, P., Brettel, K., Pignol, D., Arnoux, P., Li-Beisson, Y., Peltier, G. & Beisson, F. (2017). An algal photoenzyme converts fatty acids to hydrocarbons, Science 357, 903–907.

Sparta, K. M., Krug, M., Heinemann, U., Mueller, U. & Weiss, M. S. (2016). XDSAPP2.0, J. App. Cryst. 49, 1085–1092.

Svensson, O., Malbet-Monaco, S., Popov, A., Nurizzo, D. & Bowler, M. W. (2015). Fully automatic characterization and data collection from crystals of biological macromolecules, Acta Cryst. D 71, 1757–1767.

Tsai, Y., McPhillips, S. E., Gonzalez, A., McPhillips, T. M., Zinn, D., Cohen, A. E., Feese, M. D., Bushnell, D., Tiefenbrunn, T., Stout, C. D., Ludaescher, B., Hedman, B., Hodgson, K. O. & Soltis, S. M. (2013). AutoDrug: fully automated macromolecular crystallography workflows for fragment-based drug discovery, Acta Cryst. D 69, 796803.

Vijayachandran, L. S., Viola, C., Garzoni, F., Trowitzsch, S., Bieniossek, C., Chaillet, M., Schaffitzel, C., Busso, D., Romier, C., Poterszman, A., Richmond, T. J. & Berger, I. (2011). Robots, pipelines, polyproteins: Enabling multiprotein expression in prokaryotic and eukaryotic cells, J. Struct. Biol. 175, 198–208.

Vonrhein, C., Flensburg, C., Keller, P., Sharff, A., Smart, O., Paciorek, W., Womack, T. & Bricogne, G. (2011). Data processing and analysis with the autoPROC toolbox, Acta Cryst. D 67, 293–302.

Warne, T., Serrano-Vega, M. J., Baker, J. G., Moukhametzianov, R., Edwards, P. C., Henderson, R., Leslie, A. G., Tate, C. G. & Schertler, G. F. (2008). Structure of a β_1_-adrenergic G-protein-coupled receptor, Nature 454, 486–491.

Wasserman, S. R., Koss, J. W., Sojitra, S. T., Morisco, L. L. & Burley, S. K. (2012). Rapid-access, high-throughput synchrotron crystallography for drug discovery, Trends Pharm. Sci. 33, 261–267.

Winter, G. (2010). xia2: an expert system for macromolecular crystallography data reduction, J. App. Cryst. 43, 186–190.

Zander, U., Cianci, M., Foos, N., Silva, C. S., Mazzei, L., Zubieta, C., de Maria, A. & Nanao, M. H. (2016). Merging of synchrotron serial crystallographic data by a genetic algorithm, Acta Cryst. D 72, 1026–1035.

Zander, U., Hoffmann, G., Cornaciu, I., Marquette, J. P., Papp, G., Landret, C., Seroul, G., Sinoir, J., Rower, M., Felisaz, F., Rodriguez-Puente, S., Mariaule, V., Murphy, P., Mathieu, M., Cipriani, F. & Marquez, J. A. (2016). Automated harvesting and processing of protein crystals through laser photoablation, Acta Cryst. D 72, 454–466.

